# Cholinergic Interneurons Drive Motivation by Promoting Dopamine Release in the Nucleus Accumbens

**DOI:** 10.1101/2022.11.06.515335

**Authors:** Ali Mohebi, Val L. Collins, Joshua D. Berke

## Abstract

Motivation to work for potential rewards is critically dependent on dopamine (DA) in the nucleus accumbens (NAc). DA release from NAc axons can be controlled by at least two distinct mechanisms: 1) action potentials propagating from DA cell bodies in the ventral tegmental area (VTA), and 2) activation of β2* nicotinic receptors by local cholinergic interneurons (CINs). How CIN activity contributes to NAc DA dynamics in behaving animals remains unclear. We monitored DA release in the NAc Core of awake, unrestrained rats while simultaneously monitoring or manipulating CIN activity at the same location. CIN stimulation rapidly evoked DA release, and in contrast to slice preparations, this DA release showed no indication of short-term depression or receptor desensitization. The sound of food delivery evoked a brief joint increase in CIN population activity and DA release, with a second joint increase as rats approached the food. In an operant task, we observed fast ramps in CIN activity during approach behaviors, either to start the trial or to collect rewards. These CIN ramps co-occurred with DA release ramps, without corresponding changes in the firing of VTA DA neurons. Finally, we examined the effects of blocking CIN influence over DA release through local NAc infusion of DH*β*E, a selective antagonist of 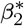 nicotinic receptors. DH*β*E dose-dependently interfered with motivated approach decisions, mimicking the effects of a DA antagonist. Our results support a key influence of CINs over motivated behavior via the local regulation of DA release.

## Introduction

Obtaining rewards typically takes work: time-consuming sequences of unrewarded actions before the reward is reached. Deciding when such work is worthwhile is an essential aspect of adaptive behavior. NAc DA is a critical modulator of such motivated decision-making (1, 2), especially choices to approach potential rewards (3, 4). Yet how NAc DA release is itself dynamically regulated to achieve appropriate motivation is not well understood.

Some features of DA release mirror the spiking of DA cell bodies. Burst firing (or strong artificial stimulation) of VTA DA cell bodies is accompanied by a corresponding pulse of NAc DA release (5–7). This pulse can encode reward prediction error (RPE), an important learning signal (8). Other aspects of DA release appear to be dissociated from spiking. Enhancing motivation by increasing reward availability does not change VTA DA cell firing rates (5, 9), but does boost NAc Core DA levels, as measured on a time scale of minutes by microdialysis (10, 11). On faster time scales (seconds), many groups have reported ramps in NAc DA as animals approach rewards (10, 12–14). These ramps may reflect discounted estimates of future reward (value), a useful motivational signal. Yet we found no evidence for corresponding ramps in spiking rate among identified DA cells in the lateral VTA (5), which provide the major DA input to NAc Core (15).

An alternative means of controlling DA release involves local circuit components within the NAc (16). Prominent among these are cholinergic interneurons (CINs), which despite constituting only 1-2% of striatal neurons, provide a very dense network of acetylcholine (ACh) fibers intermeshed with the DA axon network (17). DA axons possess nicotinic ACh receptors (nAChRs) that specifically contain the *β*_2_ subunit (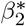; (18)). Activation of nAChRs is sufficient to locally evoke action potentials in DA axons, and consequent DA release, even in the absence of DA cell bodies (19–22).

Yet how CINs contribute to NAc DA dynamics in behaving animals is largely unknown. Slice studies suggest that DA release is evoked only by highly synchronous activation of multiple CINs (20), as might occur in response to a salient cue (23). Furthermore, DA release in slices shows a lack of summation to repetitive CIN stimulation (20), and strong paired-pulse depression (21, 24). This is apparently due to rapid desensitization of AChRs (25, 26) and feedback inhibition of CINs via DA D2 receptors (27). These features might limit the ability of CINs to sculpt motivation-related DA release, including ramping. Furthermore, results on CIN contributions to motivation are mixed. Suppression of CINs has been reported to produce a depression-like state in some behavioral tests (28) but to boost motivation in others (29).

We examined the relationships between CIN activity, DA release, and motivation. We took advantage of recent advances in optical DA sensors (30) and a specific instrumental task in which we previously found a dissociation between DA spiking and release (5). We report that CINs can indeed control DA release in awake behaving animals, but with quite distinct temporal characteristics compared to slices. We find that CIN population activity ramps up during approach behaviors in parallel with DA ramps. Furthermore, selective pharmacological blockade of NAc 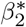 nAChRs interferes with motivated approach, as does blocking DA itself. Our results provide convergent evidence for CINs as a key mechanism regulating motivation via local control of DA release.

## Results

### ACh reliably drives DA release in NAc, without rapid depression

To study the rapid influence of CINs over striatal DA release in freely moving animals, we employed an all-optical approach. We expressed the excitatory opsin ChR2 in NAc CINs using ChAT-Cre rats (32) and Cre-dependent expression from a virus (AAV5::DIO::EF1a::ChR2::eYFP). In the same region, we expressed a red-shifted DA sensor (using AAV-DJ::CAG::RdLight1). We recorded DA dynamics by fiber photometry while driving CINs through the same fiber, in awake unrestrained rats not performing any particular task (Fig. 1a). CIN stimulation immediately increased the DA signal, and this increase was maintained as long as stimulation was applied (up to 4s; Fig. 1b, c). In contrast to results in slices (20), the amplitude of DA release scaled with the duration and frequency of laser pulses (Fig. 1d,e). CIN-evoked DA release remained robust to the second of a pair of pulses across a range of inter-pulse intervals (Fig. 1f), indicating that neither the directly evoked ACh release, nor the consequent DA release, show rapid synaptic depression in behaving animals (see also (29) for related results with choline measurements).

**Fig. 1.**
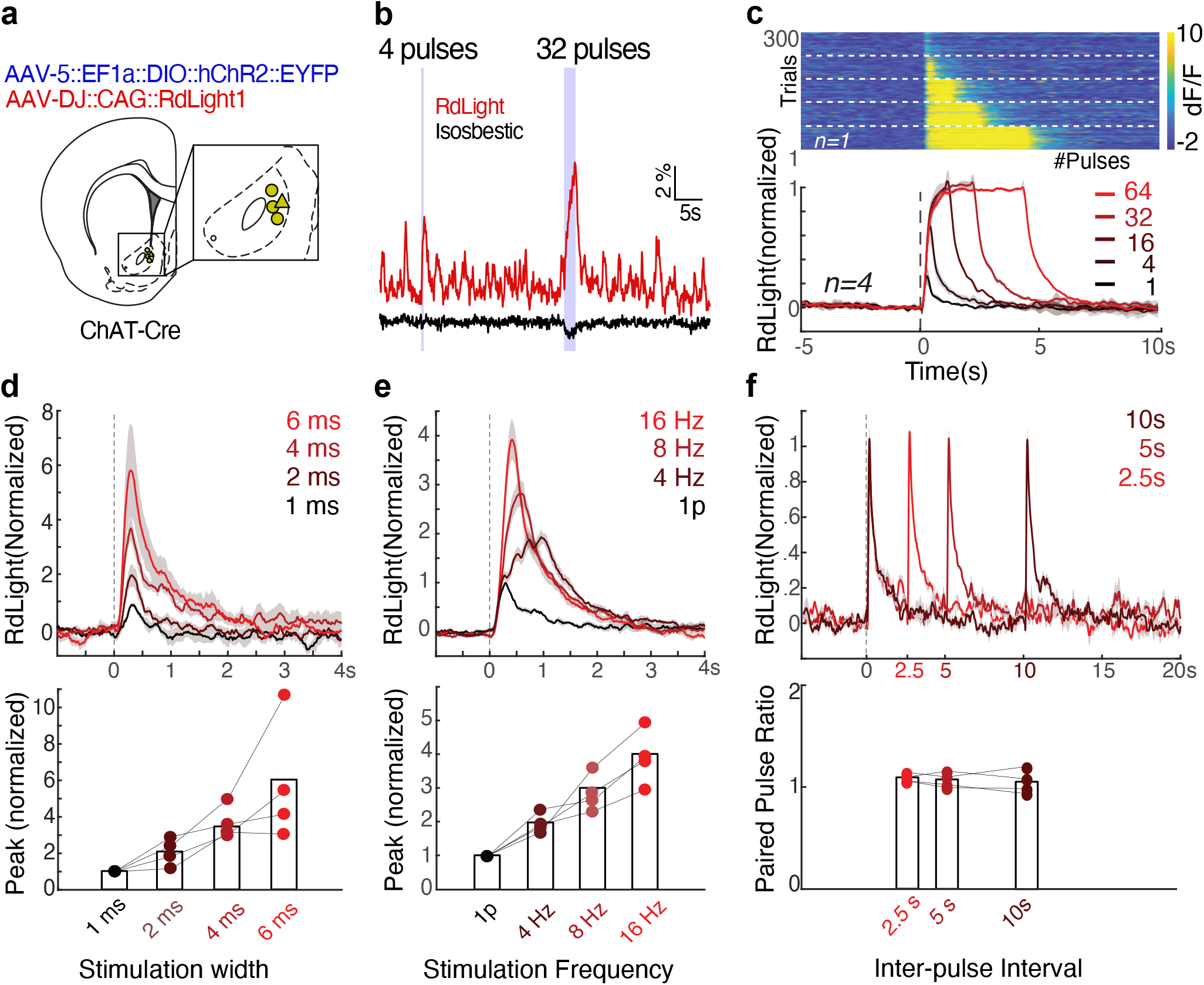
CIN stimulation drives dopamine release in freely-moving rats. **a**, Rat brain atlas (31) showing the approximate location of fiber tips. **b**, Representative traces of DA release (565 nm excitation, red) and isosbestic control (405 nm, purple), recorded from the location marked by a triangle in (a). Laser stimulation was delivered as two trains of pulses (470nm, 10mW, 4ms, 16Hz). Scale bars: 5s, 2% dF/F. **c**, *Top,* example session showing DA release in response to 16Hz optogenetic stimulation of NAc CINs, with 1,4,16,32, or 64 pulses, in pseudo-random order. Dashed lines separate trials of similar stimulation parameters. *Bottom,* normalized dF/F from RdLight aligned to the first pulse in a train of laser stimulation of NAc CINs (Stim parameters: 4 ms pulse, 16Hz, 10mW; n= 4 rats). Band shows ± SEM. dF/F traces for each recording were normalized to the maximum response evoked by 64 stimulation pulses. **d**, DA release in response to a single laser pulse of varying duration (1,2,4,6ms) at 10mW. Responses are normalized to 1ms width evoked response for each subject and averaged. Band shows ± SEM. The magnitude of DA release depends on the pulse width (ANOVA: F(3,12)=5.52,p=0.013) **e**, DA release in response to four laser pulses (4ms) of varying frequency (4,8,16 Hz; all 10mW). Release patterns are normalized to the single pulse response of the same width and power for each subject, and averaged. Band shows ± SEM. The magnitude of DA release depends on the frequency of stimulation (bottom, ANOVA: F(3,12)=9.17,p=0.002) **f**, Paired-pulse ratio test: DA release patterns in response to a pair of 4ms, 10mW pulses. A two-way ANOVA revealed that there was not a statistically significant interaction between the pulse order (first or second) and the delay between two pulses (F (2,6)=1.05, p=0.35), and there were no main effects of inter-pulse interval (p=0.47) or pulse order (p=0.06).

### CIN activity and DA release each respond to reward-related cues and ramp up during motivated approach

We next examined how CIN activity and DA release co-vary in freely moving rats. We expressed the Ca2+ indicator GCaMP6f in CINs (AAV5::Syn::Flex::GCaMP6f), and RdLight1 in the same area (AAV-DJ::CAG::RdLight1). This allowed us to monitor DA release and CIN Ca2+ dynamics through the same fiber (Fig. 2a). CIN activity and DA release showed distinct time courses (Fig. 2b), but both showed rapid increases in response to the unexpected sound of reward delivery (food hopper “Click”, occurring at random intervals drawn from a uniform distribution of 15-30s). After delivery, rats took a variable amount of time to collect the food rewards, allowing us to distinguish between neural changes associated with the sensory cue and those associated with the subsequent food retrieval. Aligning signals on food port entry revealed that both CIN GCaMP and DA release rapidly ramped up as rats approached the food port (Fig. 2 c-e).

**Fig. 2.**
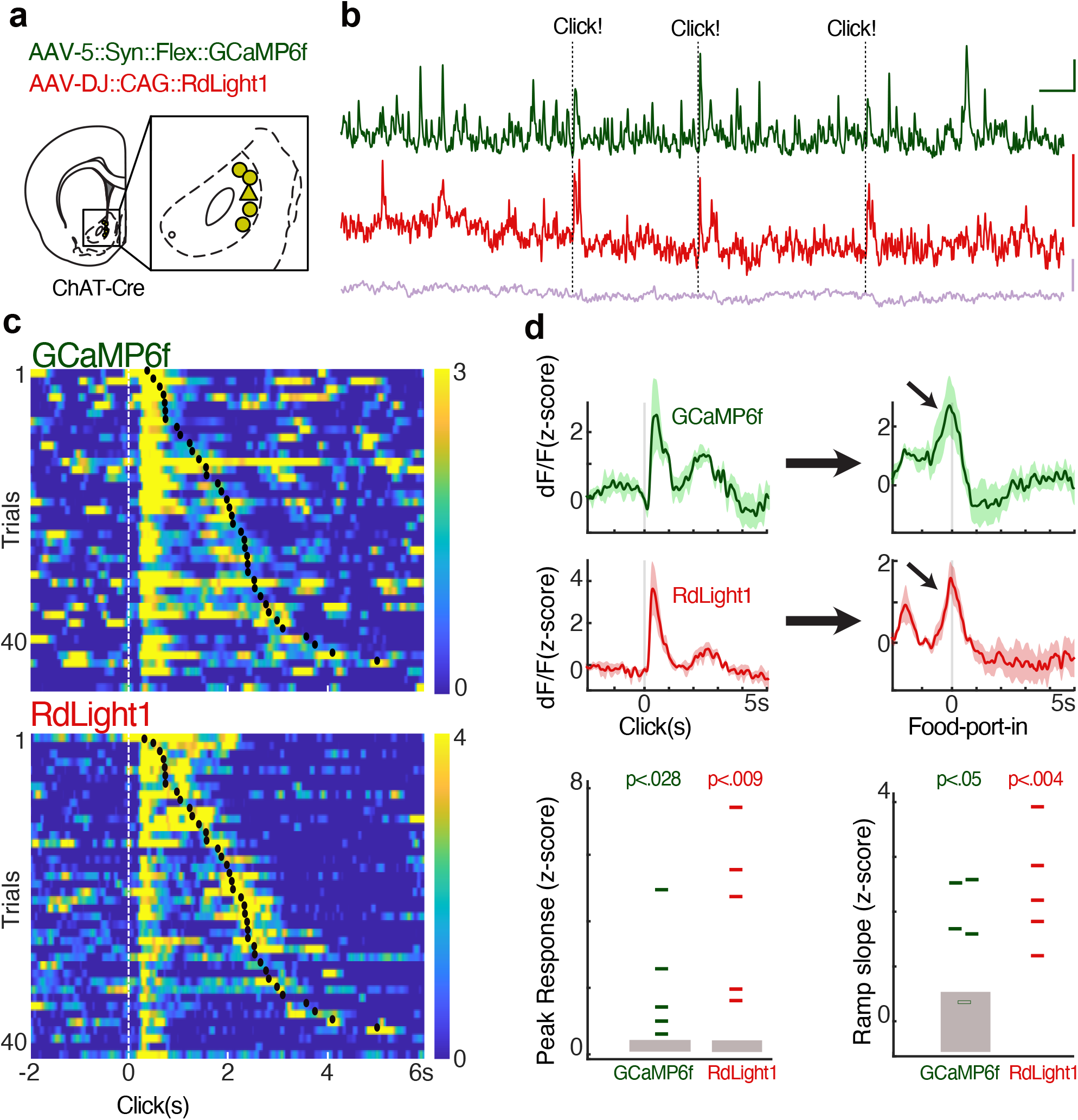
CIN activity and DA release similarly respond to reward delivery and ramp up with the reward approach. **a**, Dual-color photometry of CIN activity and DA fluctuations measured simultaneously through the same fiber. The locations of recording fiber tips are shown projected onto the nearest rat brain atlas section (31). **b**, Example of simultaneous recording of spontaneous activity, from the triangle location in (a). Green: CIN GCaMP6f signal (excitation: 470nm), red: RdLight1 (565nm), purple: isosbestic control (405nm). Dashed lines, unexpected food hopper clicks, delivering a sucrose pellet. Scale: 1% dF/F, 5s **c**, A representative session with simultaneous CIN GCaMP6f and RdLight1 recording, aligned to hopper clicks. Trials are sorted by the reward collection time (the black dot indicates food-port entry). Both CIN activity and DA fluctuations show a rapid response to the click, and a separate ramping increase during the food-port approach is apparent for longer collection times. Color scale indicates z-scored signal range. **d**, Top, average traces showing CIN activity and DA fluctuations aligned to the hopper click, peak responses (within 1s from click) significantly differed from peak values aligned to random time points in the task (n=1000 shuffles). Bottom, the slope of the signal ramps was significantly different from ramp slopes aligned to random time points throughout the session (n=1000 shuffles; grey rectangle shows 95% range). The slope was determined by fitting a line that connects the maximum and minimum signal, 0.5s before the food-port entry.

### CIN ramps can account for DA release ramps in the absence of DA firing changes

We next turned to a trial- and-error operant task, in which we have previously observed increases in NAc DA release during motivated approach without apparent increases in VTA DA firing (5). In brief, each trial starts with the illumination of a center port (Fig 3a). To obtain sucrose pellet rewards, rats approach and place their nose in this center port, and wait for a variable period (500-1500ms) until an auditory ‘Go cue’. They then nose-poke an adjacent port to the left or right. These left/right choices are probabilistically rewarded (10, 50, or 90%, probabilities change independently after blocks of 35-45 trials). Rewarded trials are made apparent by a hopper click, after which rats approach the food port to collect the pellets (Fig 3b). We examined CIN GCaMP6f signals, to assess whether ramps in CIN activity might provide the “missing” controller of DA release without changes in DA cell firing. To help disentangle signals related to sensory cue onset versus approach behavior, we examined trials in which Light-On and Center-In are more than 1s apart (i.e., “latency” > 1s). We measured ramping as the slope of the signal in the last 0.5s before approach completion. We compared these slopes to a 95% confidence interval generated by measuring slopes at random times during the task (1000 shuffles). None of the optogenetically-identified lateral VTA DA cells (0/29) showed significant ramps preceding Center-In, but ramps were reliably observed in both DA release (10/10 fiber placements in 7 rats) and CIN GCaMP (7/8 fiber placements, 5 rats). Repeating the same analysis for the Food-Port-In event produced significant ramping up for none of the 29 DA cells (one ramped down instead), but virtually all recordings of DA release (9/10) and CIN GCaMP (8/8) recordings. These observations are consistent with the hypothesis that ramps in NAc DA release are sculpted via CIN activity rather than DA cell firing.

**Fig. 3.**
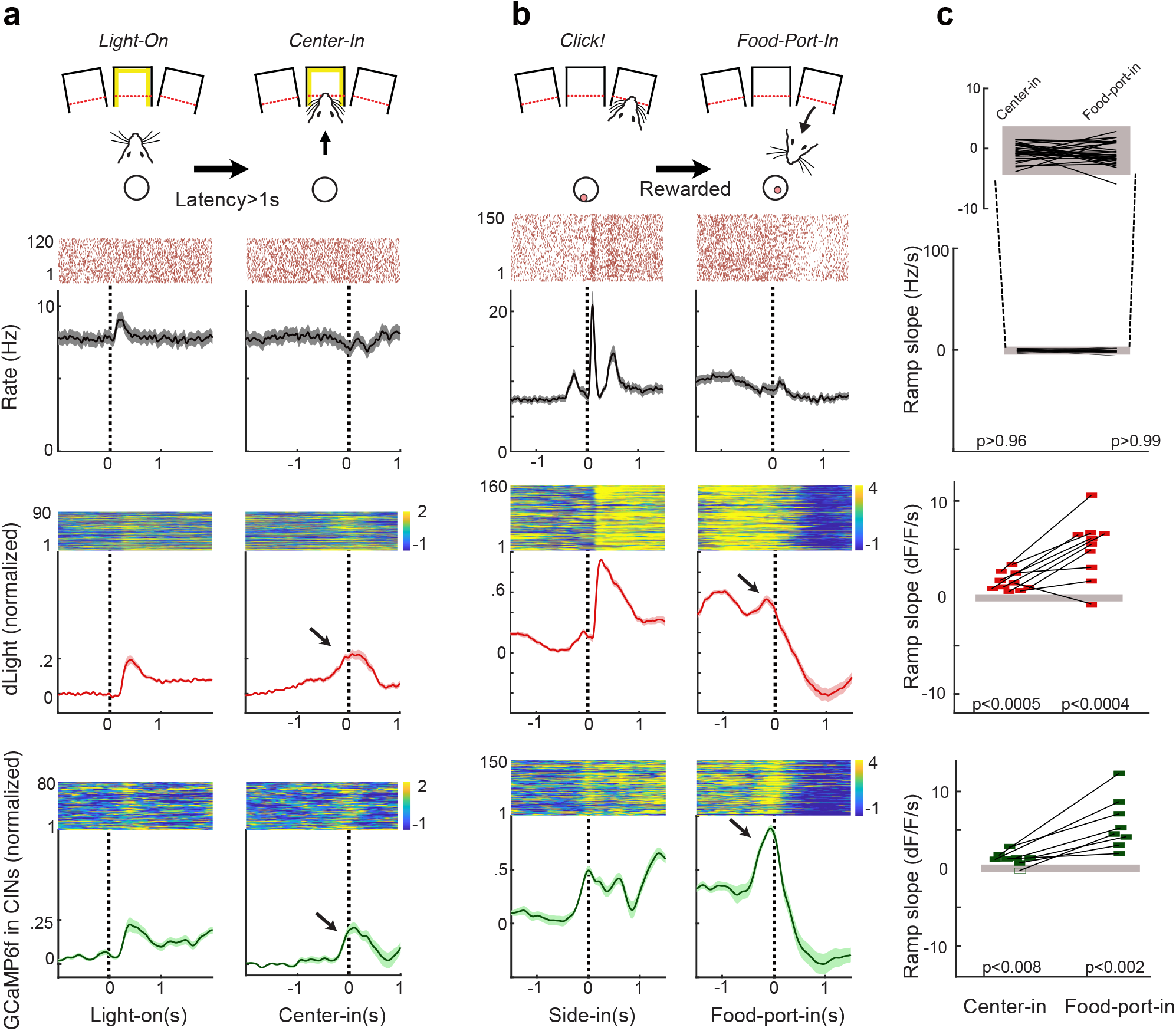
In an operant task CIN activity and DA release ramp up without corresponding increases in VTA DA cell firing. **a**, Illustration at top: trial initiation involves an approach to an illuminated center port. Data panels: Upper and middle data rows, previously-reported (5) spiking of identified VTA DA cells (n=29) and dLight DA signals (n=10). Lower row shows CIN GCaMP signals (n=8). Each panel shows a representative single example on top and the population average on the bottom. Shading indicates ± SEM. **b**, same as (a), but for the approach to the food port following the food delivery Click! sound, on rewarded trials. **c**, Quantification of slopes during approach. Data points indicate individual DA neurons (top) or fiber placements (middle, bottom). Filled grey rectangles show chance levels (95% confidence ranges, from 1000 shuffles).

### Motivated approach relies on NAc *β*_2_-containing nicotinic receptors

If NAc CINs directly act upon nearby axons to boost DA release and thereby enhance approach behaviors, such behavior should be sensitive to the blockade of the relevant nAChRs. We tested this in the trial-and-error task, using local drug infusions into the NAc Core (bilaterally, volume 0.5*μ*L / side). On consecutive days, rats received either aCSF (vehicle), DH*β*E (selective antagonist of β2* nAChRs (33, 34); 15 or 30*μ*g / side), or the non-selective dopamine antagonist flupenthixol (FLU; 10*μ*g / side), 10 minutes before beginning the task. The drug treatment order was randomly assigned and counterbalanced across rats. Blockade of 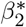 nAChRs reduced the number of completed trials in a dose-dependent manner, similarly to (though not as strongly as) FLU (Fig 4b). Both FLU and the higher dose of DH*β*E increased the latency to initiate a trial (Fig 4c). Specifically, both the higher dose of DH*β*E and FLU decreased the instantaneous likelihood (hazard rate) of decisions to approach the center port, shortly after the center port became illuminated (Fig 4d). These observations are consistent with our prior finding that bidirectional optogenetic manipulations of dopamine affect approach hazard rates during the same period (10), and support the hypothesis that nAChR-mediated DA release in NAc is involved in the motivation to work.

**Fig. 4.**
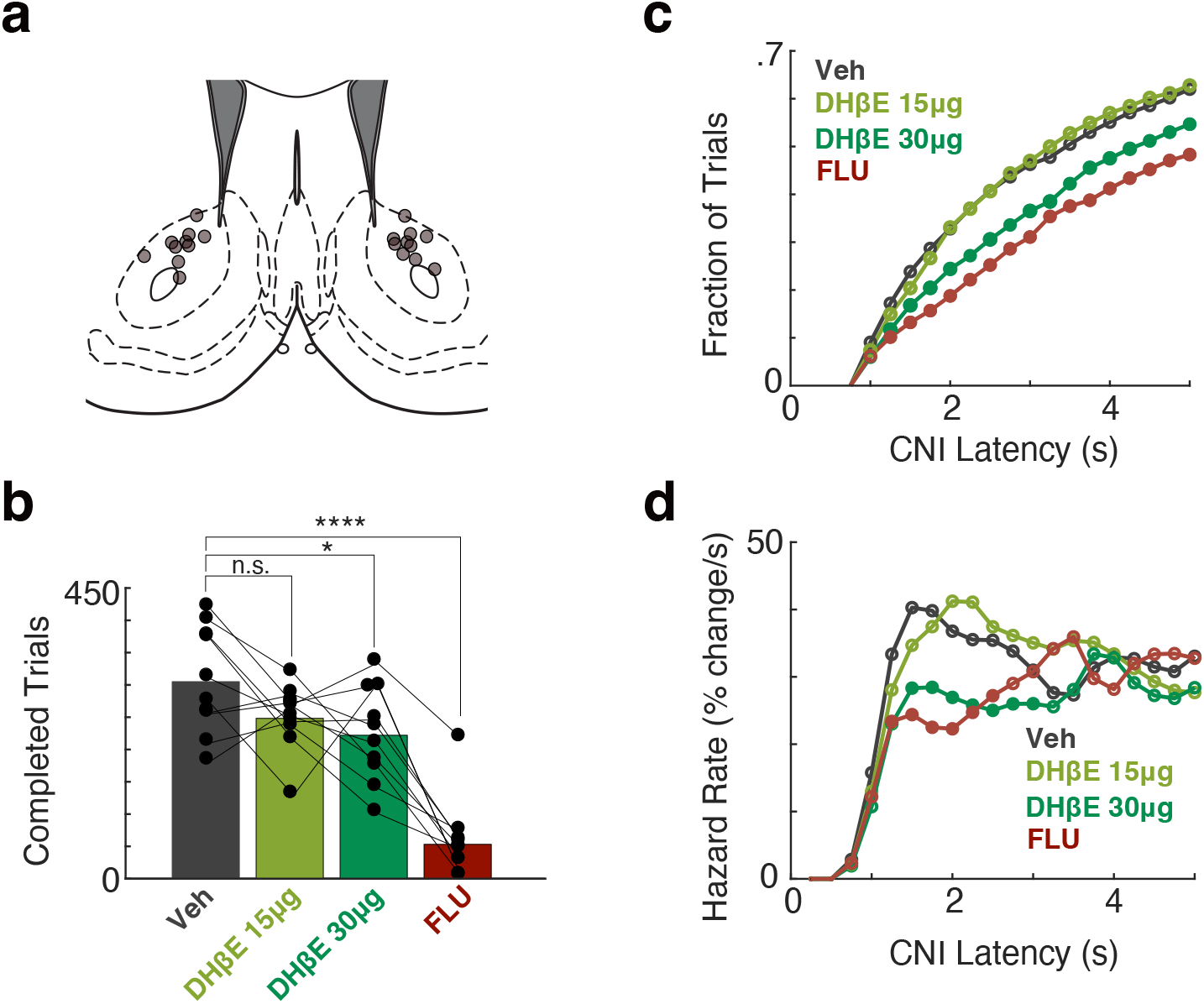
Blocking NAc dopamine or β2* nicotinic receptors diminishes motivation to work in the operant task. **a**, Placement of infusion cannula tips (circles) in NAc core, based on post-mortem histology. **b**, The number of trials completed during a 2-hr Bandit session compared to the Veh session. F(3,36)=24.64, p<7.6×10^-9^. Rats treated with DH*β*E-30*μ*g and FLU completed fewer trials compared to the vehicle session (p<0.013, p<1×10^-9^, respectively). DH*β*E-15*μ*g treatment caused a trend towards fewer completed trials (p=0.074) **c**, Cumulative distribution of long latencies (latency>1s) for different treatment conditions, binned at 250ms. Filled circles denote statistical differences from the Vehicle condition (p<0.01). **d**, The hazard rate of long latencies for different condition treatments demonstrates that blocking nicotinic transmission decreases the likelihood of a self-initiated approach (between 1-2s from light-on) in a dose-dependent manner. Filled circles denote statistical differences from the veh condition (p<0.01).

## Discussion

Local regulation of striatal DA by acetylcholine has long been established in brain slices (35). However, how this influence shapes DA signals in vivo has remained obscure. As a result, cholinergic modulation of DA release has not typically been incorporated into functional accounts of DA dynamics (Berke 2018), which have instead emphasized the faithful transmission of reward prediction errors encoded in DA cell spiking. An accurate characterization of local modulation of NAc DA in behaving animals is essential not simply for understanding the neural control of motivation but also altered physiological states evoked by disorders or abused drugs (notably, nicotine).

Seminal slice studies reported that while CINs can profoundly influence DA release via β2* nicotinic receptors, this effect shows strong short-term depression that can take minutes to recover (20, 24). If these desensitizing mechanisms apply in the awake behaving case, we might expect little functional contribution of CINs to DA signaling, and that stimulating CINs would not evoke appreciable extra DA release. Our results clearly demonstrate otherwise: stimulating CINs evoked strong DA release in a frequencydependent manner, with no evidence of depression. This result both confirms the presence of powerful ACh modulation of DA release under physiological conditions and demonstrates that this modulation has the dynamic range to be a major, continuing influence over behaviorally-relevant DA signals.

What can explain the discrepancy between ex vivo and in vivo experiments? One possibility involves the very different conditions for acetylcholinesterase (AChE), an enzyme that provides critical constraints over ACh signaling dynamics. AChE functions very rapidly, almost approaching the rate of ACh diffusion (36, 37), but this rate will be slower at the lower temperature of slice experiments compared to whole animals. Furthermore, AChE is soluble in the extracellular matrix (38), and so can be partly washed away in the slice preparation. Taken together, released ACh in the slice may escape degradation in the synaptic space for longer than normal, and thereby provoke desensitization of a larger proportion of nAChRs.

One of the most prominent reported features of striatal CINs is a reliable pause in response to reward-predicting cues, in contrast to DA release (39). However, at least in NAc we found that both DA release and CIN activity in NAc increase rapidly in response to such cues (see also (40)). Of course bulk CIN GCaMP signals are at best an incomplete readout of either CIN firing or the spatiotemporal dynamics of ACh release. In particular, imaging of CINs GCaMP in the dorsal striatum found a mismatch between signals at cell bodies and the rest of the field of view (considered to be neuropil (41)). Understanding whether control of DA release via nAChRs directly reflects CIN firing will require more experiments using either optogenetic tagging of CINs (42) or high-resolution voltage imaging (43). A related important direction for future studies is to understand why some transient fluctuations in CIN activity evoke DA release (19) but others do not (e.g. Fig. 2b).

Given that ACh promotes NAc DA release, a continuing conundrum is to explain long-standing evidence for opposing roles for striatal DA and ACh in behavioral control (29, 44, 45). One potentially-relevant behavioral dimension is whether actions are prompted by the onset of discrete external cues, versus being “self-initiated”. Increases in both presumed-CIN activity and striatal DA release have been reported in conjunction with self-timed actions (e.g. (46, 47)). This seems consistent with our observation of prominent joint DA and CIN activity ramps in longer-latency trials in our operant task, where rats are not producing a simple motor reaction to the trial start light, but rather approaching at a variable time after its onset. We therefore expected that DHβE would suppress the hazard rate of approach across a wide range of delays after the light onset. In fact, both DHβE and FLU particularly depressed approach behavior in the 1-3s range after light onset, with the hazard rate returning to control levels by 4s (Fig. 4d). This suggests that the NAc joint ACh-DA mechanisms investigated here are more involved in deciding whether it is worthwhile reorganizing ongoing behavior to respond to a cue (48).

Finally, if CINs do indeed help locally sculpt motivational aspects of DA release, a natural question becomes how relevant CIN signals are themselves generated. CINs receive inputs from a wide range of areas that can supply motivational information and influence DA release, including the basolateral amygdala (49), parafascicular thalamus (50), frontal cortex (51), and long-range GABAergic subpopulations in the VTA (52, 53). They also sample the state of surrounding networks of striatal projection cells, allowing them to potentially extract useful decision variables (54). Resolving how motivational signals are computed remains an ongoing challenge, but monitoring and manipulating each of these various circuit components within the same behavioral context is a promising approach.

## ACKNOWLEDGEMENTS

We thank members of the Berke Laboratory for their comments on a prior version of the manuscript. Funding was provided by the National Institute on Drug Abuse (R01DA045783), the National Institute of Neurological Disorders and Stroke (R01NS123516, R01NS116626), the National Institute on Alcohol Abuse and Alcoholism (R21AA027157), the National Institute on Mental Health (K01MH126223), Brain and Behavior Research Foundation (BBRF, NARSAD YIA 29361), and the UCSF Wheeler Center for the Neurobiology of Addiction.

## AUTHOR CONTRIBUTIONS

A.M. performed the behavioral photometry experiments and data analysis. V.C. performed the pharmacological manipulation experiment. J.B. oversaw the study. A.M. and J.B. wrote the manuscript together.

## Methods

### Behavioral Task

All animal procedures were approved by the University of California, San Francisco Animal Care Committee. Rats were trained in the operant “bandit” task described in detail previously (5, 10). Reward probabilities for left and right choices were pseudorandomly chosen from 10 50, or 90% for blocks of 35-45 trials, over a 2-hr session. Procedural errors (e.g. failure to hold until the Go cue) resulted in illumination of the house light and a time-out period of 1.5s.

### Photometry and optogenetic stimulation

We used a viral approach to express the genetically-encoded optical DA sensor RdLight1 and the Ca2+ indicator GCaMP6f. Under isoflurane anesthesia, a 1 μl cocktail of AAV-DJ-CAG-RdLight1 and AAV5-Syn-Flex-GCaMP6f was slowly (100 nl/min) injected (Nanoject III, Drummond) through a glass micropipette targeting nucleus accumbens: (AP: 1.7, ML: 1.7, DV: 7.0 mm relative to bregma). During the same surgery, optical fibers (400 μm core, 430 μm total diameter) attached to a metal ferrule (Doric) were inserted (target depth 200 μm higher than AAV) and cemented in place. Data were collected >3 weeks later, to allow for transgene expression. We used time-division multiplexing (55): green (565 nm), blue (470 nm) and violet (405 nm; isosbestic control) LEDs were alternately switched on and off in 10ms frames (4ms on, 6ms off). Excitation power at the fiber tip was set to 30 μW for each wavelength. Both excitation and emission signals passed through minicube filters (Doric) and bulk fluorescence was measured with a femtowatt detector (Newport, Model 2151) sampling at 10 kHz. The separate signals were then rescaled to each other via a least-square fit before analysis.

To combine fiber photometry with optogenetics, we infused a cocktail of AAV-DJ-CAG-RdLight1 and AAV5-EF1a-DIO-ChR2-eYFP into the nucleus accumbens and implanted 400μm fibers above the infusion site during the same surgery. We used time division multiplexing with interleaved LED illumination (565nm, 405nm), as above. We used brief laser pulses (470nm, 20mW) to activate ChR2.

### Drugs and Infusion Procedures

Rats were bilaterally infused with either the selective *α*_4_*β*_2_ nicotinic antagonist DH*β*E (low dose: 15*μ*g/.5*μ*L/side, high dose 30*μ*g/.5*μ*L/side; Tocris Bioscience, Bristol, UK) the non-selective dopamine receptor antagonist Flupenthixol (15μg/.5μl/side; Tocris) or sterile saline (.5*μ*L/side), which served as the vehicle control. All rats received all drug infusions, counterbalanced for order. Drugs were infused as described previously (56). Briefly, dummies were removed, and injectors were inserted into the guide cannulae protruding 3mm past the guide tip targeted to the NAc. Using a micro infusion pump (Harvard Apparatus), drugs were infused at a rate of 5.*μ*L/min for 1 min. Following a 1min post-infusion wait time, injectors were removed and dummies inserted. Rats were given 10min post-infusion before being placed in operant chambers for the bandit task.

